# Scaling tree-based automated machine learning to biomedical big data with a dataset selector

**DOI:** 10.1101/502484

**Authors:** Trang T. Le, Weixuan Fu, Jason H. Moore

**Author notes:** Funded by National Institute of Health Grant Nos. LM010098, LM012601. These authors contributed equally to this work (Trang T. Le, Weixuan Fu). Direct correspondence to.

## Abstract

Automated machine learning (AutoML) systems are helpful data science assistants designed to scan data for novel features, select appropriate supervised learning models and optimize their parameters. For this purpose, Tree-based Pipeline Optimization Tool (TPOT) was developed using strongly typed genetic programming to recommend an optimized analysis pipeline for the data scientist’s prediction problem. However, TPOT may reach computational resource limits when working on big data such as whole-genome expression data. We introduce two new features implemented in TPOT that helps increase the system’s scalability: Dataset selector and Template. Dataset selector (DS) provides the option to specify subsets of the features as separate datasets, assuming the signals come from one or more of these specific data subsets. Built in at the beginning of each pipeline structure, DS reduces the computational expense of TPOT to only evaluate on a smaller subset of data rather than the entire dataset. Consequently, DS increases TPOT’s efficiency in application on big data by slicing the dataset into smaller sets of features and allowing genetic programming to select the best subset in the final pipeline. Template enforces type constraints with strongly typed genetic programming and enables the incorporation of DS at the beginning of each pipeline. We show that DS and Template help reduce TPOT computation time and may provide more interpretable results. Our simulations show TPOT-DS significantly outperforms a tuned XGBoost model and standard TPOT implementation. We apply TPOT-DS to real RNA-Seq data from a study of major depressive disorder. Independent of the previous study that identified significant association with depression severity of the enrichment scores of two modules, in an automated fashion, TPOT-DS corroborates that one of the modules is largely predictive of the clinical diagnosis of each individual.

**Author Summary:** Big data have recently become prevalent in many fields including meteorology, complex physics simulations, large scale imaging, genomics, biomedical research, environmental research and more. TPOT is a Python Automated Machine Learning (AutoML) tool that uses genetic programming to optimize machine learning pipelines for analyzing biomedical data. However, like other AutoML tools, when analyzing big data, the early implementations of TPOT face the challenges of long runtime, high computational expense as well complex pipeline with low interpretability. Here, we develop two novel features for TPOT, Dataset Selector and Template, that leverage domain knowledge, greatly reduce the computational expense and flexibly extend TPOT’s application to biomedical big data analysis.

## Introduction

For many bioinformatics problems of classifying individuals into clinical categories from highdimensional biological data, performance of a machine learning (ML) model depends greatly on the problem it is applied to [1,2]. In addition, choosing a classifier is merely one step of the arduous process that leads to predictions. To detect patterns among features (*e.g*., clinical variables) and their associations with the outcome (*e.g*., clinical diagnosis), a data scientist typically has to design and test different complex machine learning (ML) frameworks that consist of data exploration, feature engineering, model selection and prediction. Automated machine learning (AutoML) systems were developed to automate this challenging and time-consuming process. These intelligent systems increase the accessibility and scalability of various machine learning applications by efficiently solving an optimization problem to discover pipelines that yield satisfactory outcomes, such as prediction accuracy. Consequently, AutoML allows data scientists to focus their effort in applying their expertise in other important research components such as developing meaningful hypotheses or communicating the results.

Various approaches have been employed to build AutoML systems for diverse applications. Auto-sklearn [3] and Auto-WEKA [4] use Bayesian optimization for model selection and hyperparameter optimization. Recipe [6] optimizes the ML pipeline through grammar-based genetic programming and Autostacker [7] automates stacked ensembling. Both methods automate hyperparameter tuning and model selection using evolutionary algorithm. DEvol [8] designs deep neural network specifically via genetic programming. H2O.ai [9] automates data preprocessing, hyperparameter tuning, random grid search and stacked ensembles in a distributed ML platform in multiple languages. Finally, Xcessiv [10] provides web-based application for quick, scalable, and automated hyper-parameter tuning and stacked ensembling in Python.

Tree-based Pipeline Optimization Tool (TPOT) is a genetic programming-based AutoML system that uses genetic programming (GP) [11] to optimize a series of feature selectors, preprocessors and ML models with the objective of maximizing classification accuracy. While most AutoML systems primarily focus on model selection and hyperparameter optimization, TPOT also pays attention to feature selection and feature engineering by evaluating the complete pipelines based on their cross-validated score such as mean squared error or balanced accuracy. Given no a priori knowledge about the problem, TPOT has been shown to frequently outperform standard machine learning analyses [12,13]. Effort has been made to specialize TPOT for human genetics research, resulting in a useful extended version of TPOT, TPOT-MDR, that features Multifactor Dimensionality Reduction and an Expert Knowledge Filter [14]. However, at the current stage, TPOT still requires great computational expense to analyze large datasets such as in genome-wide association studies (GWAS) or gene expression analyses. Consequently, the application of TPOT on real-world datasets has been limited to small sets of features [15].

In this work, we introduce two new features implemented in TPOT that helps increase the system’s scalability. First, the Dataset Selector (DS) allows the users to pass specific subsets of the features, reducing the computational expense of TPOT at the beginning of each pipeline to only evaluate on a smaller subset of data rather than the entire dataset. Consequently, DS increases TPOT’s efficiency in application on large data sets by slicing the data into smaller sets of features (*e.g*. genes) and allowing a genetic algorithm to select the best subset in the final pipeline. Second, Template enables the option for strongly typed GP, a method to enforce type constraints in genetic programming. By letting users specify a desired structure of the resulting machine learning pipeline, Template helps reduce TPOT computation time and potentially provide more interpretable results.

## Methods

We begin with descriptions of the two novel additions to TPOT, Dataset Selector and Template. Then, we provide detail of a real-world RNA-Seq expression dataset and describe a simulation approach to generate data comparable to the expression data. Finally, we discuss other methods and performance metrics for comparison. Detailed simulation and analysis code needed to reproduce the results has been made available on the GitHub repository https://github.com/trang1618/tpot-ds.

### Tree-based Pipeline Optimization Tool

Tree-based Pipeline Optimization Tool (TPOT) automates the laborious process of designing a ML pipeline by representing pipelines as binary expression trees with ML operators as primitives. Pipeline elements include algorithms from the extensive library of scikit-learn [16] as well as other efficient implementations such as extreme gradient boosting. Applying GP with the NSGA-II Pareto optimization [17], TPOT optimizes the accuracy achieved by the pipeline while accounting for its complexity. Specifically, to automatically generate and optimize these machine learning pipelines, TPOT utilizes the Python package DEAP [18] to implement the GP algorithm. Implementation details can be found at TPOT’s active Github repository https://github.com/EpistasisLab/tpot.

### Dataset Selector

TPOT’s current operators include sets of feature pre-processors, feature transformers, feature selection techniques, and supervised classifiers and regressions. In this study, we introduce a new operator called Dataset Selector (DS) that enables biologically guided group-level feature selection. Specifically, taking place at the very first stage of the pipeline, DS passes only a specific subset of the features onwards, effectively slicing the large original dataset into smaller ones. Hence, with DS, users can specify subsets of features of interest to reduce the feature space’s dimension at pipeline initialization. From predefined subsets of features, the DS operator allows TPOT to select the best subset that maximizes average accuracy in *k*-fold cross validation (5-fold by default).

For example, in a gene expression analysis of major depressive disorder, a neuroscientist can specify collections of genes in pathways of interest and identify the important collection that helps predict the depression severity. Similarly, in a genome-wide association study of breast cancer, an analyst may assign variants in the data to different subsets of potentially related variants and detect the subset associated with the breast cancer diagnosis. In general, the DS operator allows for compartmentalization of the feature space to smaller subsets based on *a priori* expert knowledge about the biomedical dataset. From here, TPOT learns and selects the most relevant group of features for outcome prediction.

### Template

Parallel with the establishment of the Dataset Selector operator, we now offer TPOT users the option to define a Template that provides a way to specify a desired structure for the resulting machine learning pipeline, which will reduce TPOT computation time and potentially provide more interpretable results.

Current implementation of Template supports linear pipelines, or path graphs, which are trees with two nodes (operators) of vertex degree 1, and the other *n* − 2 nodes of vertex degree 2. Further, Template takes advantage of the strongly typed genetic programming framework that enforces data-type constraints [19] and imposes type-based restrictions on which element (*i.e*., operator) type can be chosen at each node. In strongly typed genetic programming, while the fitness function and parameters remain the same, the initialization procedure and genetic operators (*e.g*., mutation, crossover) must respect the enhanced legality constraints [19]. With a Template defined, each node in the tree pipeline is assigned one of the five major operator types: dataset selector, feature selection, feature transform, classifier or regressor. Moreover, besides the major operator types, each node can also be assigned more specifically as a method of an operator, such as decision trees for classifier. An example Template is Dataset selector → Feature transform → Decision trees.

### Datasets

We apply TPOT with the new DS operator on both simulated datasets and a real world RNA-Seq gene expression dataset. With both real-world and simulated data, we hope to acquire a comprehensive view of the strengths and limitations of TPOT in the next generation sequencing domain.

#### Simulation methods

The simulated datasets were generated using the `R` package `privateEC`, which was designed to simulate realistic effects to be expected in gene expression or resting-state fMRI data. In the current study, to be consistent with the real expression dataset (described below), we simulate interaction effect data with *m* = 200 individuals (100 cases and 100 controls) and *p* = 5,000 real-valued features with 4% functional (true positive association with outcome) for each training and testing set. Full details of the simulation approach can be found in Refs. [20,21]. Briefly, the privateEC simulation induces a differential co-expression network of random normal expression levels and permutes the values of targeted features within the cases to generate interactions. Further, by imposing a large number of background features (no association with outcome), we seek to assess TPOT-DS’s performance in accommodating large numbers of non-predictive features.

To closely resemble the module size distribution in the RNA-Seq data, we first fit a Γ distribution to the observed module sizes then sample from this distribution values for the simulated subset size, before the total number of features reaches 4,800 (number of background features). Then, the background features were randomly placed in each subset corresponding to its size. Also, for each subset *S_i_, i* = 1, …, *n*, a functional feature *s_j_* belongs to the subset with the probability

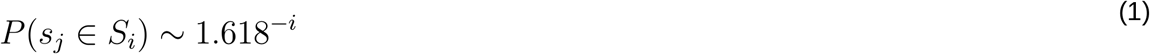

where 1.618 is an approximation of the golden ratio and yields a reasonable distribution of the functional features: they are more likely to be included in the earlier subsets (subset 1 and 2) than the later ones.

#### Real-world RNA-Seq expression data

We employed TPOT-DS on an RNA-Seq expression dataset of 78 individuals with major depressive disorder (MDD) and 79 healthy controls (HC) from Ref. [20]. Gene expression levels were quantified from reads of 19,968 annotated protein-coding genes and underwent a series of preprocessing steps including low read-count and outlier removal, technical and batch effect adjustment, and coefficient of variation filtering. Consequently, whole blood RNA-Seq measurements of 5,912 genes were obtained and are now used in the current study to test for association with MDD status. We use the 23 subsets of interconnected genes called depression gene modules (DGMs) identified from the RNA-Seq gene network module analysis [20] as input for the DS operator. We remark that these modules were constructed by an unsupervised machine learning method with dynamic tree cutting from a co-expression network. As a result, this prior knowledge of the gene structure does not depend on the diagnostic phenotype and thus yields no bias in the downstream analysis of TPOT-DS.

### Performance assessment

For each simulated and real-world dataset, after randomly splitting the entire data in two balanced smaller sets (75% training and 25% holdout), we trained TPOT-DS with the Template `Dataset Selector-Transformer-Classifier` on training data to predict class (*e.g*., diagnostic phenotype in real-world data) in the holdout set. We assess the performance of TPOT-DS by quantifying its ability to correctly select the most important subset (containing most functional features) in 100 replicates of TPOT runs on simulated data with known underlying truth. To prevent potential overfitting, we select the pipeline that is closest to the 90th percentile of the crossvalidation accuracy to be optimal. This rationale is motivated by a similar procedure for optimizing the penalty coefficient in regularized regression where the most parsimonious model within one standard error of the minimum cross-validation error is picked []. We compare the holdout (out-of-sample) accuracy of TPOT-DS’s optimal pipeline on the holdout set with that of standard TPOT (with `Transformer-Classifier` Template, no DS operator) and extreme Gradient Boosting [22], or XGBoost, which is a fast and an efficient implementation of the gradient tree boosting method that has shown much utility in many winning Kaggle solutions [23] and been successfully incorporated in several neural network architectures [24,25]. In the family of gradient boosted decision trees, XGBoost accounts for complex non-linear interaction structure among features and leverages gradient descents and boosting (sequential ensemble of weak classifiers) to effectively produce a strong prediction model. To obtain the optimal performance for this baseline model, we tune XGBoost hyperparameters using TPOT Template with only one classifier `XGBClassifier`, which is imported from the `xgboost` python package. Because of stochasticity in the optimal pipeline from TPOT-DS, standard TPOT and the tuned XGBoost model, we fit these models on the training data 100 times and compare 100 holdout accuracy values from each method. We choose accuracy to be the metric for comparison because phenotype is balanced in both simulated data and real-world data.

### Manuscript drafting

This manuscript is collaboratively written using Manubot [26], a software that supports open paper writing via GitHub using the Markdown language. Manubot uses continuous integration to monitor changes and automatically update the manuscript. Consequently, the latest version of this manuscript is always available at https://trang1618.github.io/tpot-ds-ms/.

## Results

Our main goal is to test the performance of methods to identify features that discriminate between groups and optimize the classification accuracy.

### The general workflow of TPOT-DS

The template `Dataset Selector-Transformer-Classifier` specifies a simple general workflow of TPOT-DS (Fig. 1).

**Figure 1:**
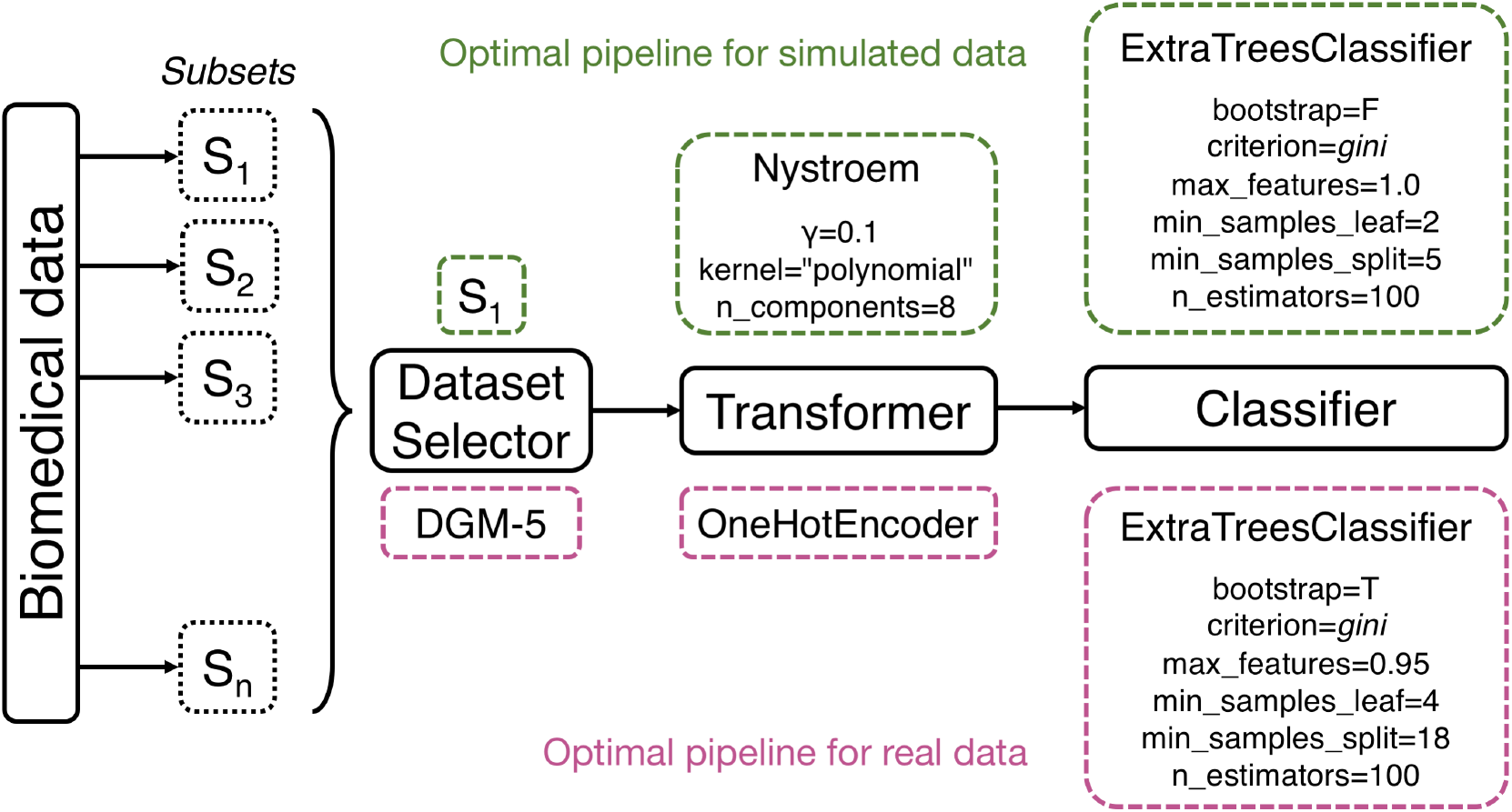
TPOT-DS’s workflow and example pipelines. Final pipelines with optimized parameters are shown for simulated data (top) and real-world gene expression data (bottom).

As discussed earlier in the Methods section, the optimal pipeline from TPOT-DS and standard TPOT is selected to be closest to the 90th percentile of the cross-validation accuracy. The optimal model of XGBoost holds properly tuned hyperparameters. For simulated dataset, the optimal pipeline selects subset *S*_1_ then constructs an approximate feature map for a linear kernel with Nystroem, which uses a subset of the data as the basis for the approximation. The final prediction is made with an extra-trees classifier that fits a number of randomized decision trees on various sub-samples of the dataset with the presented optimized parameters (Fig. 1). For the real-world dataset, the most optimal pipeline selects subset DGM-5, one-hot encode the features, then, similar to simulated data, makes the final prediction with an extra-trees classifier with a different set of optimized parameters (Fig. 1).

### Accuracy assessment of optimal pipelines

We compare the accuracy produced by optimal models from TPOT-DS, standard TPOT and XGBoost on classifying a simulated dataset with moderate interaction effect. We assign values of the effect size in the simulations to generate adequately challenging datasets so that the methods’ accuracies stay moderate and do not cluster around 0.5 or 1. The data set is split into 75% training and 25% holdout. The three models are built from the training dataset, then the trained model is applied to the independent holdout data to obtain the holdout accuracy.

We also apply the three methods to the RNA-Seq study of 78 major depressive disorder (MDD) subjects and 79 healthy controls (HC) described in [20]. The dataset contains 5,912 genes after preprocessing and filtering (see Methods for more detail). We excluded 277 genes that did not belong to 23 subsets of interconnected genes (DGMs) so that the dataset remains the same across the three methods. As with simulated data, all models are built from the training dataset (61 HC, 56 MDD), then the trained model is applied to the independent holdout data (18 HC, 22 MDD).

For the simulated data, across all 100 model fits, the optimal TPOT-DS pipeline yields an average holdout prediction accuracy of 0.65, while the standard TPOT without DS and tuned XGBoost models respectively report an average holdout accuracy of 0.48 and 0.49 (Fig. 2). This overfitting in the performance of these other two models is likely due to the models’ high flexibility that *over-learns* the training data, especially with the presence of many noisy background features.

**Figure 2:**
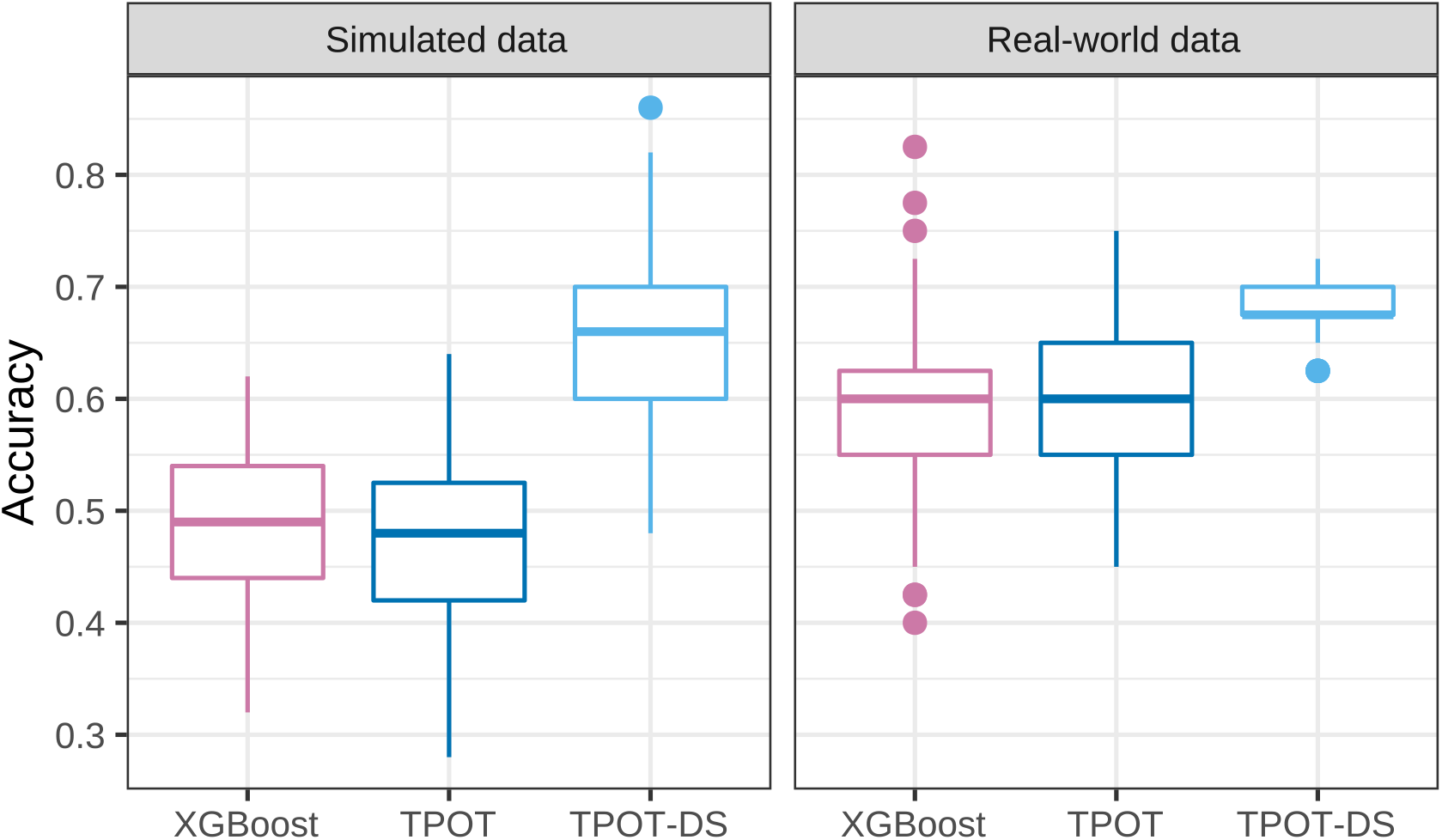
Performance comparison of three models: tuned XGBoost, optimal pipeline from standard TPOT and optimal pipeline from TPOT-DS.

Meanwhile, for the real-world expression data, the optimal TPOT-DS pipeline yields an average holdout prediction accuracy of 0.68, while the standard TPOT without DS and tuned XGBoost models produce average holdout accuracies of 0.60 and 0.59 respectively across all 100 model fits (Fig. 2). In summary, the optimal models from standard TPOT and XGBoost perform better in real-world data compared to simulated data but still worse than that of TPOT-DS. In both datasets, separate Welch two-sample one-sided *t*-tests show TPOT-DS optimal pipelines significantly outperform those of XGBoost and standard TPOT (all *p* values < 10^−15^).

Our simulation design produces a reasonable distribution of the functional features in all subsets, of which proportions are shown in Table [S1]. According to Eq. 1, the earlier the subset, the more functional features it has. Therefore, our aim is to determine how well TPOT-DS can identify the first subset (*S*_1_) that contains the largest number of informative features. In 100 replications, TPOT-DS correctly selects subset *S*_1_ in 75 resulting pipelines (Fig. 3), with the highest average holdout accuracy (0.69 across all 75 pipelines).

**Figure 3:**
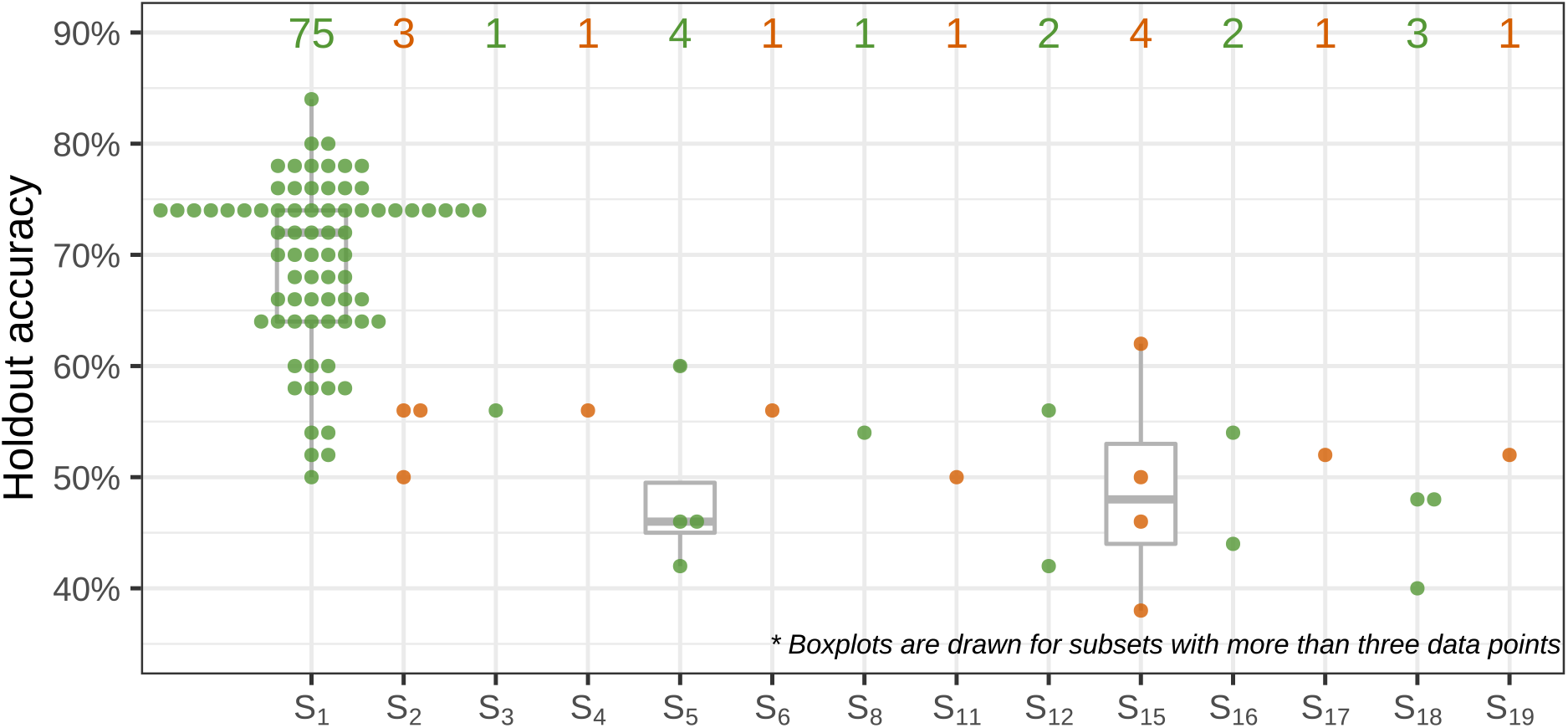
TPOT-DS’s holdout accuracy in simulated data with selected subset. Number of pipeline inclusions of each subset in 100 replications is displayed above the boxplots. Subset *S*_1_ is the most frequent to be included in the final pipeline and yields the best prediction accuracy in the holdout set.

For the expression data, in 100 replications, TPOT-DS selects DGM-5 (291 genes) 64 times to be the subset most predictive of the diagnosis status (Fig. 4), with the highest average holdout accuracy of 0.636 across 64 pipelines. In the previous study with a modular network approach, we showed that DGM-5 has statistically significant associations with depression severity measured by the Montgomery-Åsberg Depression Scale (MADRS). Although there is no direct link between the top genes of the module (Fig. 5a) and MDD in the literature, many of these genes interact with other MDD-related genes. For example, NR2C2 interacts with FKBP5 gene whose association with MDD has been strongly suggested [27,28,29]. Many of DGM-5’s top genes, including FAM13A, NR2C2,PP7080 and OXR1, were previously shown to have significant association with the diagnosis phenotype using a Relief-based feature selection method [30]. Further, with 82% overlap of DGM-5’s genes in a separate dataset from the RNA-Seq study by Mostafavi et al. [31], this gene collection’s enrichment score was also shown to be significantly associated with the diagnosis status in this independent dataset.

**Figure 4:**
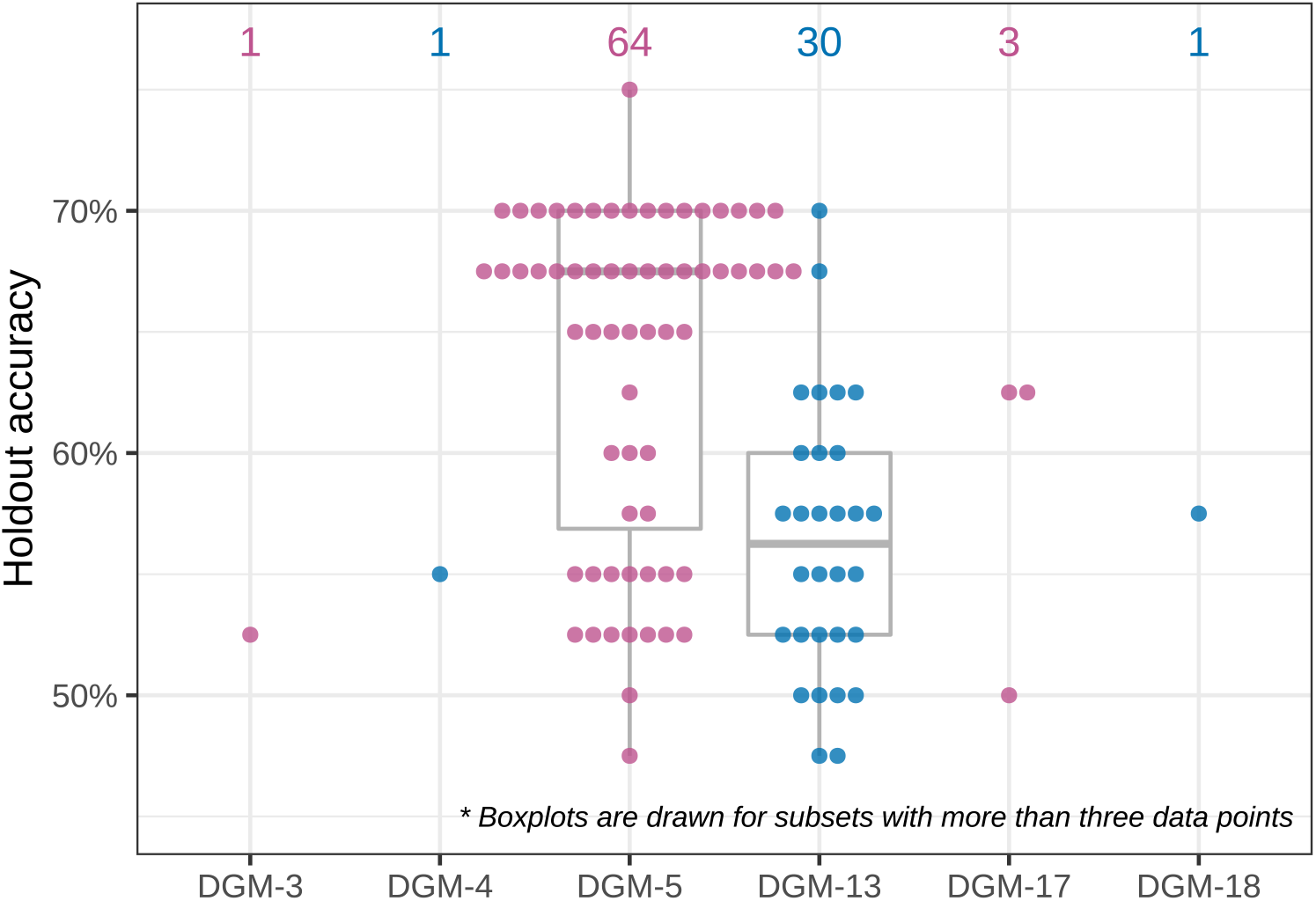
TPOT-DS’s holdout accuracy in RNA-Seq expression data with selected subset. Number of pipeline inclusions of each subset in 100 replications is displayed above the boxplots. Subsets DGM-5 and DGM-13 are the most frequent to be included in the final pipeline. Pipelines that include DGM-5, on average, produce higher MDD prediction accuracies in the holdout set.

**Figure 5:**
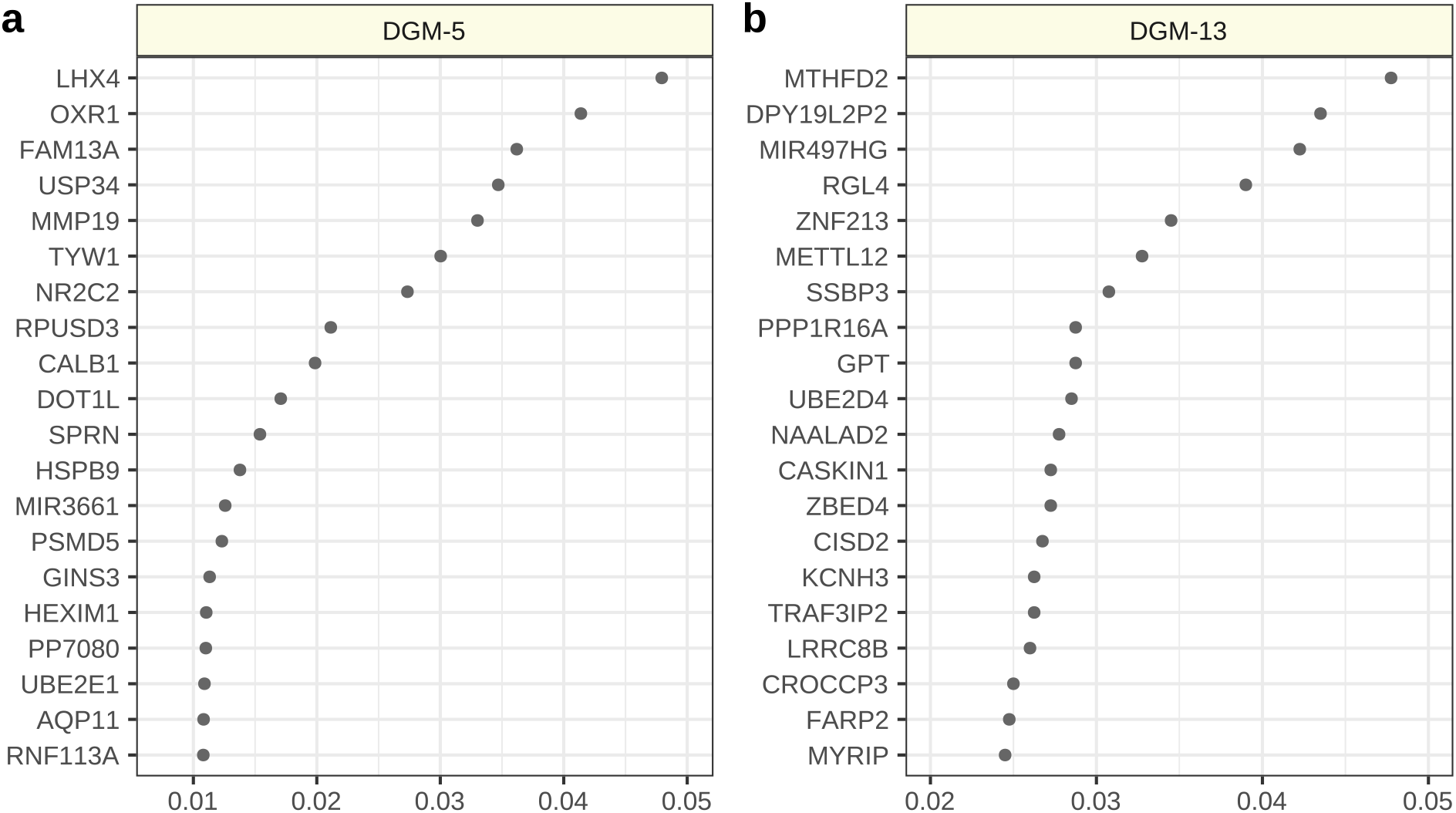
Permutation importance scores of the top twenty expression features in the optimal pipeline that selects DGM-5 and one that selects DGM-13. Comprehensive importance scores of the all expression features computed by permutation from the optimal pipelines are provided in Table S2.

After DGM-5, DGM-13 (134 genes) was selected by TPOT-DS 30 times (Fig. 4), with an average holdout accuracy of 0.563 across 30 pipelines. The previous network approach did not find statistically significant association between this module’s enrichment score and the MADRS. While many of the top genes (Fig. 5b) do not have direct disease association, several have been linked to depression-like behavior in animal studies such as PPP1R16A [32] and CASKIN1 [33]. The RGL4 gene, a Ral guanine nucleotide dissociation stimulator, was found to have a rare protein disruptive variant in at least one suicide patient among 60 other mutations [34].

### Computational expense

For a dataset of the size simulated in our study (*m* = 200 samples and *p* = 5000 attributes), standard TPOT has a 18.5-hour runtime on a low performance computing machine with an Intel Xeon E5-2690 2.60GHz CPU, 28 cores and 256GB of RAM, whereas TPOT-DS has a 65-minute runtime, approximately 17 times faster. On the same low performance computing machine, each replication of standard TPOT on the expression data takes on average 13.3 hours, whereas TPOT-DS takes 40 minutes, approximately 20 times faster.

## Discussion

To our knowledge, TPOT-DS is the first AutoML tool to offer the option of feature selection at the group level. Previously, it was computationally expensive for any AutoML program to process biomedical big data. TPOT-DS is able to identify the most meaningful group of features to include in the prediction pipeline. We assess TPOT-DS’s holdout prediction accuracy compared to standard TPOT and XGBoost, another state-of-the-art machine learning method. We apply TPOT-DS to real-world expression data to demonstrate the identification of biologically relevant groups of genes.

Implemented with a strongly typed GP, Template provides more flexibility by allowing users to prespecify a particular pipeline structure based on their knowledge, which speeds up AutoML process and provides potentially more interpretable results. For example, in high-dimensional data, dimensionality reduction or feature selection algorithms are preferably included at the beginning of the pipelines via Template to identify important features and, meanwhile, reduce computation time. For datasets with categorical features, preprocessing operators for encoding those features such as one-hot encoder should be specified in the pipeline structure to improve pipelines’ performance. Template was utilized in this study to specify the DS as the first step of the pipeline, which enables the comparison between the two TPOT implementations, with and without DS.

We simulated data of the similar scale and challenging enough for the models to have similar predictive power as in the real-world RNA-Seq data. TPOT-DS correctly selects the subset with the most important features in the majority of replications and produces high average holdout accuracy of 0.69. In both simulated and RNASeq gene expression data, the final TPOT-DS pipeline outperforms that of standard TPOT and XGBoost. The low holdout accuracies of standard TPOT and XGBoost are expected because of the few signals in a high-dimenional feature space of the data. Meanwhile, TPOT-DS finds a more compact feature space to operate on, resulting in higher prediction accuracy and lower computational expense.

Interestingly enough, TPOT-DS repeatedly selects DGM-5 to include in the final pipeline. In a previous study, we showed DGM-5 and DGM-17 enrichment scores were significantly associated with depression severity [20]. We also remarked that DGM-5 contains many genes that are biologically relevant or previously associated with mood disorders [20] and its enriched pathways such as apoptosis indicates a genetic signature of MDD pertaining to shrinkage of brain region-specific volume due to cell loss [35,36]. TPOT-DS also selects DGM-13 as a potentially predictive group of features with smaller average holdout accuracy compared to DGM-5 (0.563 < 0.636). The lack of previously found association of these genes with the phenotype is likely because MDD is a complex disorder of heterogeneous etiology [37]. Hence, the clinical diagnosis is the accumulative result of coordinated variation of many genes in the module, especially ones with high importance scores. Future studies to refine and characterize genes in DGM-13 as well as DGM-5 may deploy expression quantitative trait loci (e-QTL) or interaction QTL analysis to discover disease-associated variants [38].

Complexity-interpretability trade-off is an important topic to discuss in the context of AutoML. While arbitrarily-shaped pipelines may yield predictions competitive to human-level performance, these pipelines are often too complex to be interpretable. Vice versa, a simpler pipeline with defined steps of operators may be easier to interpret but yield suboptimal prediction accuracy. Finding the balance between pipeline complexity, model interpretation and generalization remains a challenging task for AutoML application in biomedical big data. With DS, in the terminology of evolutionary algorithm, each pipeline individual of a TPOT generation during optimization holds lower complexity due to the selected subset’s lower dimension compared to that of the entire dataset. We hope that, with the complexity reduction from imposing a strongly-type GP template and DS, a small loss in dataset-specific predictive accuracy can be compensated by considerable increase in interpretability and generalizability. In this study, the resulting TPOT-DS pipelines are more interpretable with only two simple optimized operators after the DS: a transformer and a classifier. In the case of the expression analysis, these pipelines also highlight two small sets of interconnected genes that contain candidates for MDD and related disorders. Additionally, complexity reduction results in more efficient computation, which is strongly desirable in biomedical big data analysis.

A limitation of the DS analysis is the required pre-definition of subsets prior to executing TPOT-DS. While this characteristic of an intelligent system is desirable when *a prior* knowledge on the biomedical data is available, it might pose as a challenge when this knowledge is inadequate, such as when analyzing data of a brand-new disease. Nevertheless, one can perform a clustering method such as *k*-means to group features prior to performing TPOT-DS on the data. Another limitation of the current implementation of TPOT-DS is its restricted ability to select only one subset. A future design to support tree structures for Template will enable TPOT-DS to identify more than one subset that have high predictive power of the outcome. A new operator that combines the data subsets will prove useful in this design. Extensions of TPOT-DS will also involve overlapping subsets, which will require pipeline complexity reformulation beyond the total number of operators included in a pipeline. Specifically, in the case of overlapping subsets, the number of features in the selected subset(s) is expected to be an element of the complexity calculation. Extension of TPOT-DS to GWAS is straightforward. However, because of the low predictive power of variants in current GWAS, alternative metrics beside accuracy, balanced accuracy or area under the receiving operator characteristic curve will need to be designed and included in the fitness function of TPOT’s evolutionary algorithm.

In this study, we developed two new operators for TPOT, Dataset Selector and Template, to enhance its performance on high-dimensional data by simplifying the pipeline structure and reducing the computational expense. Dataset Selector helps users leverage domain knowledge to narrow down important features for further interpretation, and Template largely increases flexibility of TPOT via customizing pipeline structure. Future extension and integration of these two operators have the potential to enrich the application of AutoML on different real world biomedical problems.

